# Genomic evidence of a widespread southern distribution during the Last Glacial Maximum for two eastern North American hickory species

**DOI:** 10.1101/205716

**Authors:** Jordan B. Bemmels, Christopher W. Dick

## Abstract

**Aim:** Phylogeographic studies of temperate forest taxa often infer complex histories involving population subdivision into distinct refugia during the Last Glacial Maximum (LGM). However, temperate forests may have been broadly distributed in southeastern North America during the LGM. We investigate genome-wide genetic structure in two widespread eastern North America tree species to determine if range expansion from a contiguous area or from genetically isolated refugia better explains the postglacial history of trees and forests from this region.

**Location:** Eastern North America (ENA).

**Taxa:** Bitternut hickory (*Carya cordiformis* (Wangenh.) K.Koch) and shagbark hickory (*Carya ovata* (Mill.) K.Koch).

**Methods:** Genetic diversity and differentiation indices were calculated from >1,000 nuclear SNP loci genotyped in ca. 180 individuals per species sampled across ENA. Genetic structure was investigated using principle component analysis and genetic clustering algorithms. As an additional tool for inference, areas of suitable habitat during the LGM were predicted using species distribution models (SDMs).

**Results:** Populations across all latitudes showed similar levels of genetic diversity. Most genetic variation was weakly differentiated across ENA, with the exception of an outlier population of *Carya ovata* in Texas. Genetic structure in each species exhibited an isolation-by-distance pattern. SDMs predicted high LGM habitat suitability over much of the southeastern United States.

**Main conclusions:** Both hickory species likely survived the LGM in a large region of continuous habitat and recolonized northern areas in a single expanding front that encountered few migration barriers. More complex scenarios, such as forest refugia, need not be invoked to explain genetic structure. The genetically distinct Texas population of *Carya ovata* could represent a separate glacial refugium, but other explanations are possible. Relative to that of other temperate forest regions, the phylogeographic history of ENA may have been exceptionally simple, involving a northward range shift but without well defined refugia.

## Introduction

Temperate forest ecosystems have long served as models for understanding how historical forces give rise to population genetic structure in terrestrial organisms (Hewitt, 1999, 2000; Petit et al., 2003; Soltis et al., 2006; Shafer et al., 2010; Qiu et al., 2011; Gavin et al., 2014; Lumibao et al., 2017). Migrational responses to Pleistocene glaciation often left large impacts on the genetic structure of temperate species, but the severity and effects of glaciation varied in different regions of the world (Hewitt, 2000; Shafer et al., 2010; Qiu et al., 2011; Lumibao et al., 2017). In Europe, where many classic phylogeographic paradigms were first established (Lumibao et al., 2017) and where glaciation during the Last Glacial Maximum (LGM, ca. 21.5 ka; Jackson et al., 2000) was particularly severe, widespread temperate taxa typically retreated to distinct glacial refugia in Mediterranean regions (Hewitt, 1999, 2000; Petit et al., 2003). Genetic diversity was progressively lost due to founder effects as populations recolonized northern areas following glacial retreat (Hewitt, 1999, 2000), except in mid-latitude areas where admixture of lineages from distinct refugia has often led to elevated genetic diversity (Petit et al., 2003).

In other temperate regions of the world, phylogeographic patterns were structured by very different geographies and glacial histories. In eastern North America (ENA), early studies tended to emphasize genetic breaks between populations separated by rivers and mountain ranges (Soltis et al., 2006; Jaramillo-Correa et al., 2009). In western North America, major refugia existed in the Pacific Northwest and Beringia, with smaller refugia on offshore islands and between continental ice sheets Shafer et al., 2010). In East Asia, responses to glaciation included not only latitudinal migration, but also elevational and longitudinal migration and *in situ* persistence (Qiu et al., 2011). The complexity of these classic paradigms has recently been expanded in all four northern hemisphere regions to include small, low-density cryptic refugia in areas previously thought unsuitable for habitation by temperate species (Stewart & Lister, 2001; Willis & Van Andel, 2004; McLachlan et al., 2005; Soltis et al., 2006; Provan & Bennett, 2008; Qiu et al., 2011). However, the molecular and fossil evidence supporting the existence of cryptic refugia is not universally accepted (Tzedakis et al., 2013).

Compared to the other three northern hemisphere temperate forest regions, ENA is phylogeographically unique for at least three reasons. Firstly, its geography is relatively simple, characterized by a large contiguous landmass with only a single north-south mountain range of modest height (i.e., the Appalachians), and generally gradual transitions between ecosystem types. Secondly, latitudinal temperature gradients during the LGM were particularly steep, with warm areas located in close proximity to glaciers (Tzedakis et al., 2013). Thirdly, despite numerous phylogeographic studies, well-delineated glacial refugia generally shared by most species have not conclusively been identified. Proposed refugial locations include the Gulf Coast, the Atlantic Coast, Florida, Texas, the Ozark Plateau, the Lower Mississippi River Valley, the Appalachians, and interior areas near ice sheets (Griffin & Barrett, 2004; Magni et al., 2005; Soltis et al., 2006; Jaramillo-Correa et al., 2009; Morris et al., 2010; Barnard-Kubow et al., 2015; McCarthy & Mason-Gamer, 2016; Peterson & Graves, 2016), which together sum to nearly the entire unglaciated region of eastern North America. While some species may have survived in one or more of these distinct refugia, fossil and genetic evidence suggests that some temperate taxa were widespread over vast areas of the southeastern United States during the LGM (Bennett, 1985; Magni et al., 2005; McLachlan et al., 2005; Peterson & Graves, 2016; Lumibao et al., 2017).

Given the diversity of hypotheses that have been evoked to explain phylogeographic patterns in ENA taxa, studies assessing genome-wide patterns of genetic variation in widely distributed model species would provide valuable insight into the history of temperate forests from the region as a whole. Surprisingly, we are aware of no such studies (but Eckert et al. (2010) and Nadeau et al. (2015) have conducted studies of more narrowly distributed tree species). Here, we use genome-wide genetic variation to examine the phylogeographic history of two widespread, ENA tree species: bitternut hickory (*Carya cordiformis* (Wagenh.) K.Koch) and shagbark hickory (*Carya ovata* (Mill.) K.Koch). We construct and analyze single-nucleotide polymorphism (SNP) datasets from nearly rangewide collections of each species and build paleodistribution models in order to characterize geographic patterns of genetic diversity and differentiation across ENA. In particular, we aim to determine if genetic structure is best explained by recolonization from distinct forest refugia (and if so, where these refugia where located), or by a single expansion from a large, continuous forest region.

## Methods

### Study species

*Carya cordiformis* and *Carya ovata* are wind-pollinated, animal-dispersed trees co-distributed from southern Quebec to eastern Texas (Fig. 1). Their ranges roughly correspond to the overall geographic distribution of temperate deciduous forests in ENA, making them excellent model taxa to study the phylogeography of eastern deciduous forests as a whole. *Carya ovata* additionally occurs in several small, disjunct populations in the Sierra Madre Oriental of northern Mexico (Little, 1971). In the northern half of temperate ENA, *C. cordiformis* occupies many habitats but occurs most frequently on mesic soils and bottomlands (Smith, 1990), whereas *C. ovata* is common on a wider variety of sites (Graney, 1990). In southern areas both species are less common and typically restricted to wet, fertile soils (Graney, 1990; Smith, 1990).

**Figure 1.**
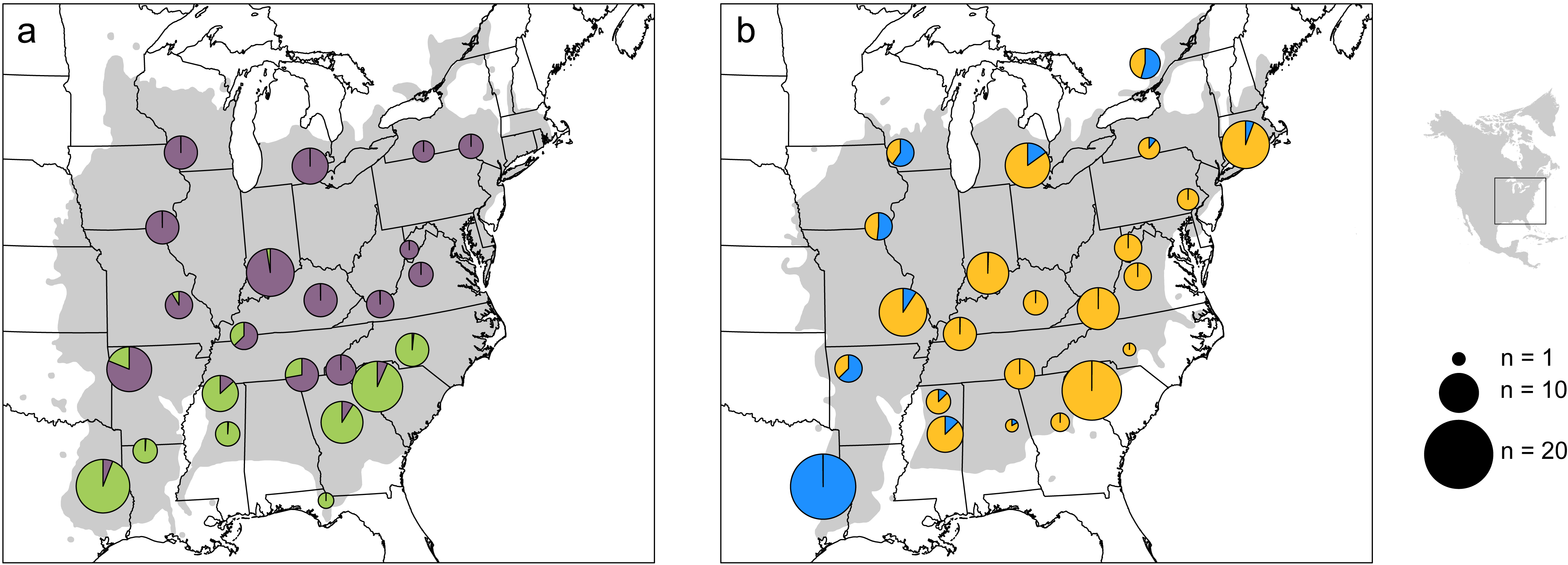
Membership of (a) *Carya cordiformis* and (b) *Carya ovata* populations in genetic clusters (different colours on pie charts; *K* = 2) identified using fastStructure (Raj et al., 2014). The geographic distribution of each species is shown in grey (Little, 1971), and the sample size (n) of each population is proportional to the size of the pie chart. Note that optimal *K* = 1-2, but *K* = 1 is not shown as all individuals of each species would belong to the same genetic cluster.

Phylogeographic knowledge is completely lacking in *C. cordiformis*. In *C. ovata*, analysis of cpDNA haplotypes has revealed no clear pattern, as some haplotypes are widespread throughout the entire range and others are more spatially restricted, including in formerly glaciated areas (Lumibao et al., 2017). *Carya* pollen is not typically distinguished to the species level, but LGM-age pollen of *Carya* has been found at low density over large areas of the southeastern USA (Prentice et al., 1991; Jackson et al., 2000). *Carya* macrofossils dating to the LGM have been found in southern Tennessee (35ºN; Jackson et al., 2000), and trace amounts of pollen as far north as southern Illinois (39ºN; Grüger, 1972). However, the presence of small amounts of pollen in northern areas is not necessarily indicative of local presence, as pollen can be reworked from layers representing different time periods, or transported by wind over long distances (Tzedakis et al., 2013).

Many of the ca. 13 North American *Carya* species readily hybridize with one another (Fralish & Franklin, 2002). Geographically structured hybridization may impact phylogeographic inferences in tree species (Saeki et al., 2011; Thomson et al., 2015), and one limitation of our analyses is that we are unable to assess patterns of hybridization with other *Carya*. However, no stable hybrid zones exist in our species and we consider it unlikely that occasional hybridization would systematically bias genetic structure in a similar way across thousands of loci.

### DNA sampling and SNP genotyping

Silica-dried leaf tissue was collected from 182 individuals of each species from populations across ENA (Fig. 1; Tables S1.1-S1.2). Sampled individuals within each population were separated by a minimum of 50 m (but sometimes up to dozens of km) in order to minimize the chance of sampling siblings and other close relatives. Sample size varied greatly among populations depending on the number of individuals meeting these requirements that could be located (mean N = 7.8; Tables S1.1-S1.2). A representative voucher specimen from each population was deposited in the University of Michigan Herbarium (MICH; DOI: XXXXX).

DNA samples were extracted using Nucleospin Plant II extraction kits (*Macherey-Nagel*; Düren, Germany), and libraries were prepared using a modified double digest Restriction Associated DNA (ddRAD) sequencing protocol following Peterson et al., (2012), with restriction enzymes *EcoRI* and *MseI.* Full details of extraction methods and library preparation are provided in Appendix S1 in Supporting Information. Seven libraries of 72 samples each were sequenced at The Hospital for Sick Children (Toronto, ON) on an *Illumina HiSeq* (Illumina; San Diego, CA) using single-end 50-bp sequencing. In order to ensure adequate depth of coverage, at least one million raw reads per sample were required to process a sample, and individuals not meeting this target were resequenced in subsequent libraries.

Loci were identified and single nucleotide polymorphisms (SNPs) were genotyped using STACKS v.1.44-v.1.46 (Catchen et al., 2011, 2013). Full details of SNP discovery are provided in Appendix S1. After SNPs were successfully identified, one SNP genotype per locus was exported from STACKS using the *populations* tool, retaining only SNPs with a minimum genotyping rate of 75% (-r 0.75) and a minimum minor allele frequency (MAF) of 3.3% (--min_maf 0.033), the lowest detectable MAF in at least one population of each species, following Massatti & Knowles (2014). Minimum MAF is an important parameter to consider because it can impact inference of genetic structure (De la Cruz & Raska, 2014). We therefore explored preliminary analyses with minimum MAF = 1% and 5%, but found that using the higher minimum MAF (5%) made little qualitative difference in preliminary results. With the lower minimum MAF (1%), broad-scale patterns were overwhelmed by local-scale signatures, likely due to rare variants shared among closely related individuals (De la Cruz & Raska, 2014). These results suggested that our choice of minimum MAF = 3.3% was appropriate.

In order to retain only putatively nuclear SNPs, we removed any SNPs from loci that aligned with a maximum of two mismatches (-v 2) to the *Juglans regia* (Juglandaceae) chloroplast genome (Genbank accession NC_028617.1) or the *Cucurbita pepo* (Cucurbitaceae) mitochondrion genome (NC_014050.1) using BOWTIE v.1.2 (Langmead et al., 2009). Extremely variable loci were also excluded as these may represent locus assembly errors; we defined these loci as those with values of θ (Watterson, 1975) above the 95^th^ percentile, with θ calculated for each locus individually using the *R* package ‘pegas’ v.0.10 (Paradis, 2010). Individual samples with unusually high levels of missing data across all loci (based on visual inspection) were also excluded.

### Genetic diversity and divergence

Three genetic diversity parameters were calculated overall and for each population: observed and expected heterozygosity (*H_o_* and *H*_*e*_, respectively), and nucleotide diversity (*π*). Genetic differentiation (*F*_*ST*_) (Nei, 1987) was calculated overall and pairwise between each pair of populations. *H_o_*, *H_e_*, and *F*_*ST*_ were calculated in the *R* package ‘hierfstat’ v.0.04-22 (Goudet, 2005), while π was calculated using *populations* in STACKS v.1.46 (Catchen et al., 2011, 2013). Genetic diversity and differentiation measures are not reported for populations represented by a single individual.

### Population genetic structure

In order to test for isolation by distance (IBD), Mantel tests (Mantel, 1967) were performed to assess the relationship between population pairwise *F*_*ST*_ values and geographic distances. Principal component analysis (PCA) was used to investigate genetic relationships among individuals and populations using the *dudi.pca* function in the *R* package ‘adegenet’ v.2.0.1 (Jombart, 2008; Jombart & Ahmed, 2011). The NC_e population (Table S1.1) was excluded for *C. cordiformis* because some, but not all individuals from this population formed a distinct genetic cluster, which suggests that several closely related individuals were unintentionally sampled and the high genetic similarity between these individuals could have biased initial PCA results.

Genetic clusters were characterized using FASTSTRUCTURE v.1.0 (Raj et al., 2014), with all populations and individuals included, using the recommended procedure for detecting subtle genetic structure. Initially, the simple prior model was used and the number of clusters (*K*) was varied from 1 to 6 for each species, and *K* was selected using the *chooseK* tool in FASTSTRUCTURE. Then, FASTSTRUCTURE was rerun 100 times using the logistic prior model for the optimal value(s) of *K*, and final estimates of genetic membership of individuals in each genetic cluster were obtained as the average membership from the five runs with the highest likelihood, following Raj et al. (2014). After investigating the broadest level of structure within each dataset, we reran FASTSTRUCTURE on individual genetic clusters in order to test for substructure within clusters.

### Paleodistribution modelling

Species distribution models (SDMs) were constructed in order to predict the potential distribution of each species during the Last Glacial Maximum (LGM; 21.5 ka). Complete details of SDM construction and data sources are given in Appendix S1. Briefly, occurrence records were obtained from the US Forest Service Forest Inventory Analysis Database (O’Connell et al., 2012), while environmental variables were obtained at 2.5-arcminute resolution from the WorldClim v.1.4. (Hijmans et al., 2005) and envirem (Title & Bemmels, 2017) databases, for both current and LGM conditions. SDMs were constructed using Maxent v.3.4.1 (Phillips et al., 2004, 2006, 2017) in the *R* package ‘dismo’ (Hijmans et al., 2015), with models optimized according to best practices, following Title & Bemmels (2017).

## Results

### Genetic diversity and differentiation

The final genetic datasets for *C. cordiformis* and *C. ovata* contained 177 individuals genotyped at 1,046 SNPs, and 180 individuals genotyped at 1,018 SNPs, respectively. The overall genotyping rate for both species was 89%.

While some populations were represented by very few individuals (Tables S1.1-S1.2), very small sample sizes are typically sufficient to obtain accurate population genomic measures of genetic diversity and differentiation if calculated across thousands of SNPs (Willing et al., 2012; Nazareno et al., 2017). Furthermore, genetic diversity estimates were generally uncorrelated with population sample size (Fig. S1.1), except that a negative relationship was found between *H_o_* and sample size in *C. ovata* (R^2^ = 0.21, p = 0.035). However, as *H*_*o*_ is computed on a per-individual basis, there is no theoretical reason to expect that sample size might bias estimates of *H*_*o*_ and we suspect that this correlation is spurious.

Genetic diversity showed little variation among populations for both species (Fig. 2). A significant decline in genetic diversity with increasing latitude was not observed for any genetic diversity measure (*H*_*o*_, *H*_*e*_, *π*) for either species. Instead, a significant increase in *H_o_* with increasing latitude was observed in *C. cordiformis* (R^2^ = 0.23, p = 0.021), as was a marginally non-significant increase in *H_o_* with increasing latitude in *C. ovata* (R^2^ = 0.19, p = 0.051). Although genetic variation was fairly uniform across latitudes, far northern and far southern populations sometimes showed slightly lower values of *H*_*e*_ and *π* than typical of mid-latitude populations (Fig. 2), as expected for range-edge populations (Jaramillo-Correa et al., 2009). Among-population genetic differentiation (*F*_*ST*_) is low in both species overall (*C. cordiformis*: 0.047; *C. ovata*: 0.038), and among most pairs of populations (Tables S1.3-S1.4).

**Figure 2.**
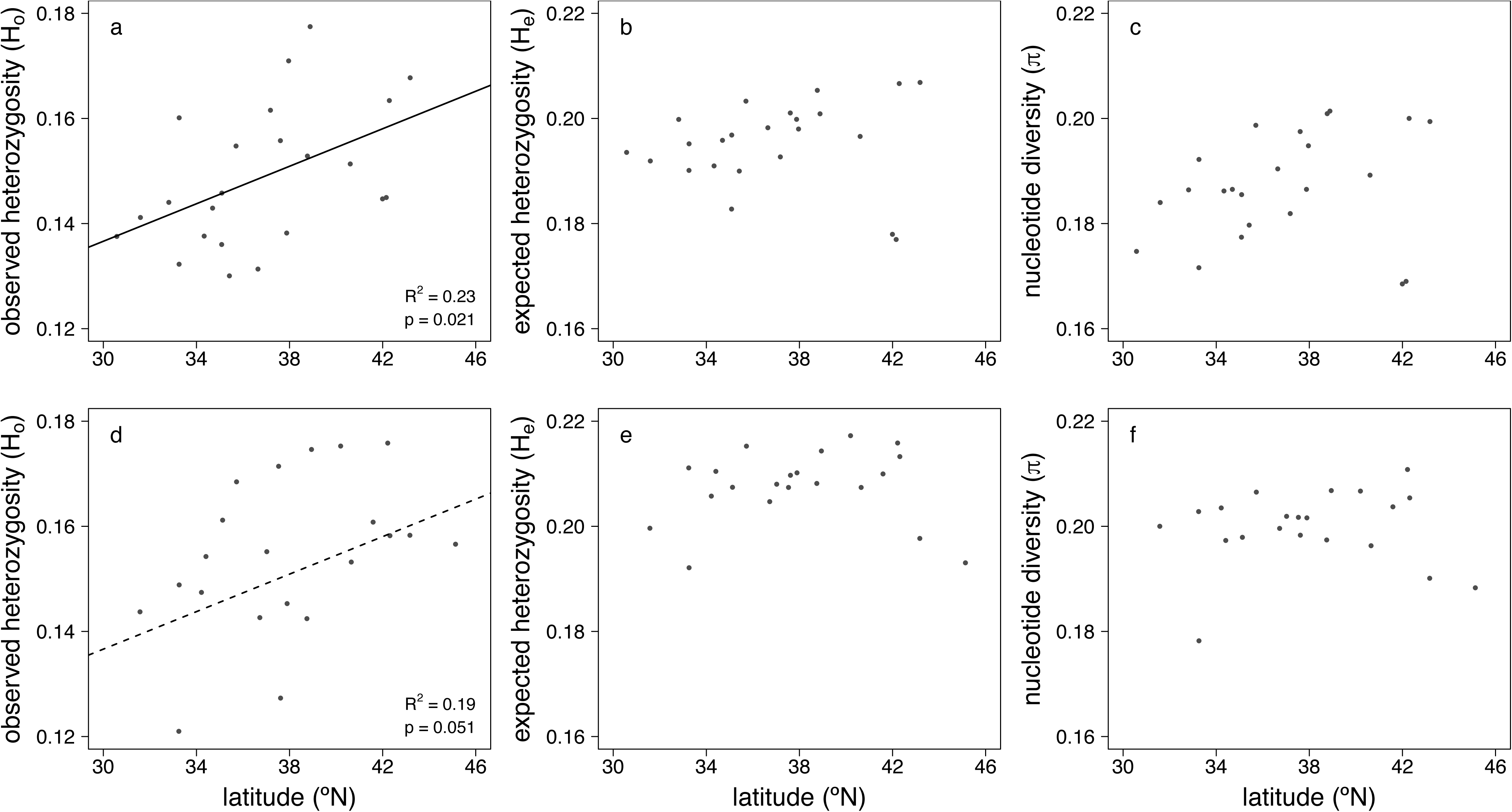
Genetic diversity vs. latitude for populations of *Carya cordiformis* (a-c) and *Carya ovata* (d-f). Statistically significant relationships are portrayed as solid lines, and marginally significant relationships as dashed lines.

### Spatial genetic structure

Spatial genetic structure in both species is very weak and dominated by a pattern of isolation by distance (IBD). Mantel tests of IBD were statistically significant in both species (*C. cordiformis*: r = 0.36, p = 0.0017; *C. ovata*: r = 0.47, p = 8.1 x 10^−5^; Fig. 3).

**Figure 3.**
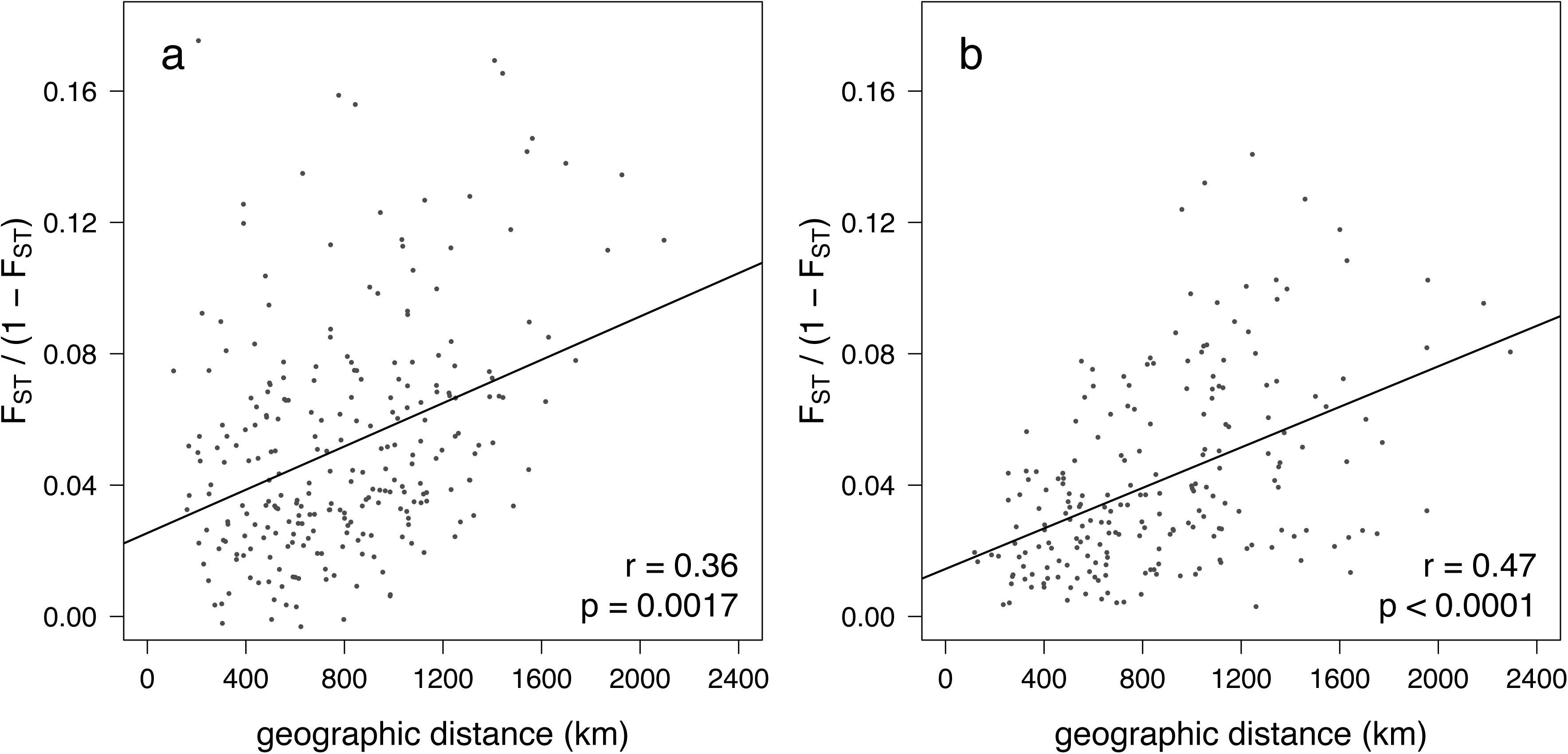
Mantel test showing isolation by distance in (a) *Carya cordiformis* and (b) *Carya ovata*. Each dot represents a pair of populations and the y-axis is a measure of genetic differentiation between populations.

Principal component analysis (PCA) also revealed an IBD-like pattern, without clearly defined, distinct genetic clusters (Fig. 4). One exception to this pattern is that in *C. ovata*, the Texas population (TX; bright pink dots, Fig. 4b) forms a separate cluster that does not overlap with any other populations. However, the relative position of TX along PC axes is still closest to that of geographically proximate western and southern populations, consistent with an IBD pattern.

**Figure 4.**
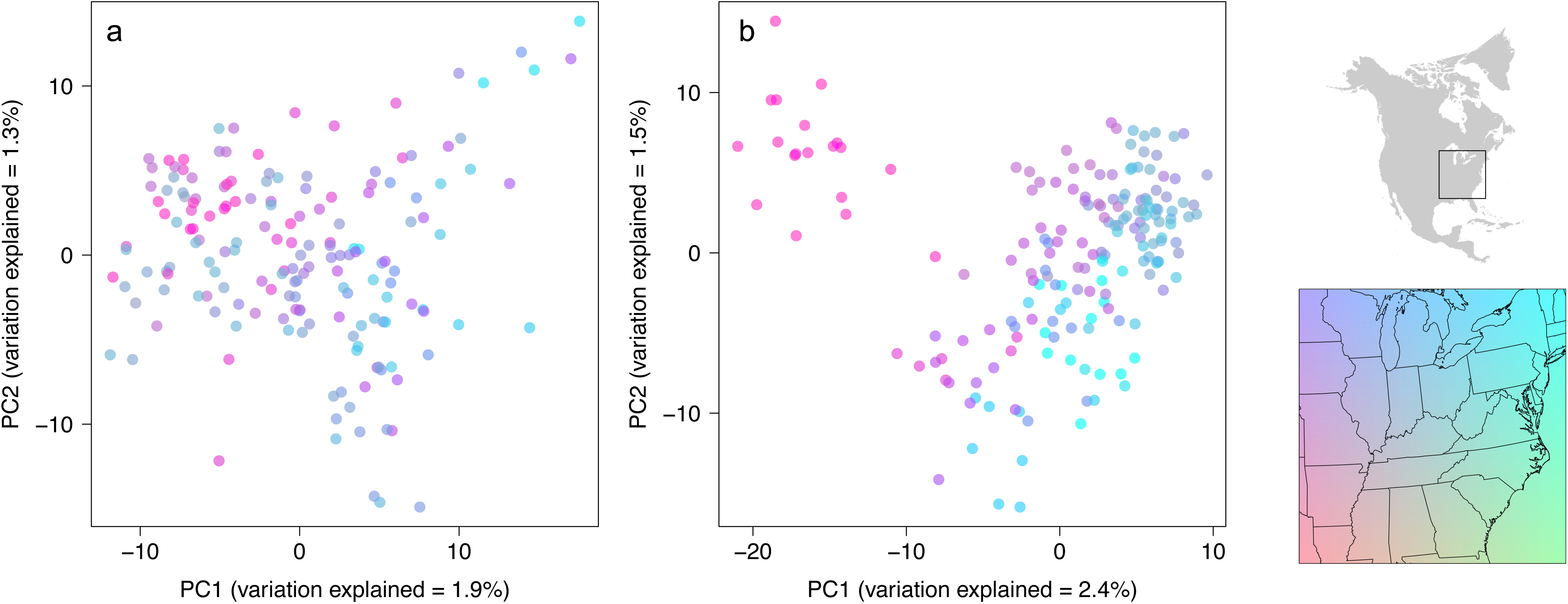
Clustering of individuals along the first and second principal component (PC) axes of genetic variation in (a) *Carya cordiformis* and (b) *Carya ovata.* Each dot represents a single individual, and colours correspond to the geographic location of the individual (shown in the inset map of eastern North America). Note that the Texas population (TX) of *C. ovata* discussed in the manuscript is represented by the pink dots in the upper left corner of (b).

Lack of strong genetic structure was also suggested by FASTSTRUCTURE results. Under the model with simple priors, the optimal number of genetic clusters was *K* = 1 for both species. Under the model with logistic priors, which is more useful for detecting subtle structure (Raj et al., 2014), optimal *K* ranged from 1-6. However, the logistic priors model is prone to overfitting (Raj et al., 2014) and *K* > 2 did not produce results that were biologically interpretable. We therefore note that *K* = 1 or 2 is likely the optimal model complexity to explain genetic structure. In both species with *K* = 2, genetic structure was weak and similar to an IBD-like pattern (Figs. 5, S1.2). In *C. cordiformis*, a gradual north-south transition between clusters was evident. In *C. ovata*, the transition between clusters was primarily from east to west. No substructure was evident within any genetic cluster for any species, except that within the western cluster for *C. ovata*, optimal *K* = 2 and the Texas population (TX) forms a distinct subcluster relative to the other four populations (AR, IA, ON, WI; data not shown). However, the grouping of these four populations into a distinct subcluster may be only a statistical artifact reflecting the substantial additional membership of each these four western populations in the main eastern cluster.

### Paleodistribution modelling

The same four climatic variables were coincidentally retained in the SDMs for both species: maximum temperature of the coldest month, potential evapotranspiration of the warmest quarter, mean annual precipitation, and climatic moisture index (Table S1.5). For both species, models were able to predict the current species distribution (Fig. 1) very well along the northern and western range edges, but performed more poorly at delineating the southern range edge (Fig. 5). This poorer performance may reflect the fact that both species are rare in the southern portion of their ranges, where presence and absence may be determined more by soil type and topography (Graney, 1990; Smith, 1990) than by broad-scale climatic differences among sites. For both species, a large, continuous area of high LGM habitat suitability is predicted to have extended over much of the southeastern US, from central Texas to exposed continental shelf off the coast of North Carolina, and south to northern Florida (Fig. 5).

**Figure 5.**
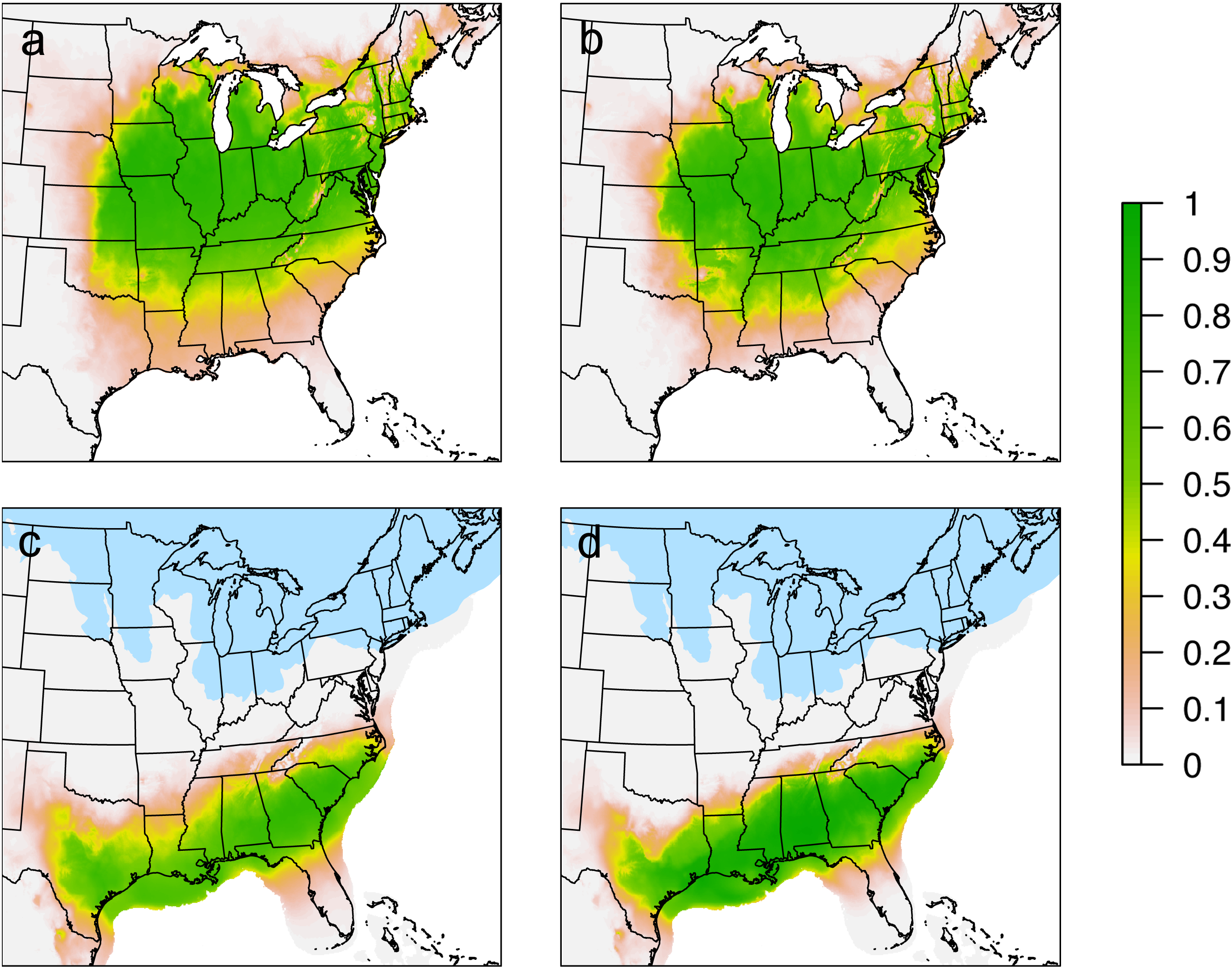
Species distribution models showing predicted habitat suitability (grey to green colour scale) for (a, c) *Carya cordiformis* and (b, d) *Carya ovata*, in both (a-b) the current time period and (c-d) the Last Glacial Maximum (ca. 21.5. Glaciated areas are shown in blue.

## Discussion

*Carya cordiformis* and *Carya ovata* likely survived the LGM in a large, contiguous forest region covering much of the southeastern United States, and recolonized northern areas in a single expanding front. This scenario is supported by our genetic results showing weak genetic structure and an isolation-by-distance (IBD) pattern, and by our paleodistribution models and the fossil record. Genetic differentiation is continuous across ENA, except that a Texas population of *C. ovata* is more genetically distinct than other populations. The cause of this distinctiveness remains unclear. Insights from both species are likely applicable to understanding the phylogeographic history of temperate forests from ENA as a whole, and suggest that ENA may lack the complex refugial dynamics characteristic of other temperate forest regions of the world.

### Weak genetic structure

In both species, genetic structure is weak and geographic patterns of genetic variation are primarily characterized by IBD, rather than sharp genetic breaks among regions. Although some geographic structure was detected with FASTSTRUCTURE, IBD is known to bias tests of hierarchical genetic structure (Frantz et al., 2009; Meirmans, 2012). In particular, such tests are susceptible to incorrect inference of multiple genetic clusters when populations are geographically subsampled from within a single larger cluster subject to IBD (Frantz et al., 2009; Meirmans, 2012). We note that a rangewide IBD pattern is present in our datasets (Fig. 3), and genetic structure is weak in both species (optimal *K* = 1 to 2), with only gradual geographic transitions between inferred genetic clusters (Fig. 1). Similarly, substantial overlap among populations exists along the first and second principal component axes (Fig. 4), with the exception of the Texas population of *C. ovata* (see below). Rather than indicating the true presence of biologically meaningful genetic clusters, our fastSTRUCTURE results are likely a statistical artifact of underlying IBD and suggest that over the majority of the species range, genetic differentiation is continuous.

In addition to lack of phylogeographic breaks, there are no identifiable regions of elevated genetic diversity (Fig. 2). Similar to many other temperate tree species from ENA (Lumibao et al., 2017), *C. cordiformis* and *C. ovata* do not exhibit declines in genetic diversity with increasing latitude, and population genetic diversity (*H*_*e*_, *π*) is relatively uniform across the species range. The absence of regions of elevated diversity suggests either historical recolonization in a single, slowly expanding migration front experiencing little loss of diversity during migration, or else very high gene flow among populations (Jaramillo-Correa et al., 2009). Both of these situations are likely applicable to our study species. In particular, low population genetic differentiation (*C. cordiformis*, *F*_*ST*_ = 0.047; *C. ovata*, *F*_*ST*_ = 0.038) provides further evidence that gene flow among populations is likely high. Low population differentiation is commonly observed in widespread, wind-pollinated forest trees, due to their large population sizes and capacity for long-distance pollen-mediated gene flow (Hamrick et al., 1992; Savolainen et al., 2007; Alberto et al., 2013).

Despite generally uniform genetic diversity across populations in terms of *H*_*e*_ and *π*, observed heterozygosity (*H_o_*) increases with increasing latitude in both species (Fig. 2). Higher genetic diversity in northern regions might reflect the effects of larger populations in the north that are less subject to genetic drift (Griffin & Barrett, 2004), but genetic drift would be expected to simultaneously affect *H*_*e*_ and *π* (not only *H*_*o*_). A more likely explanation is that decreased *H*_*o*_ in the south reflects an increase in homozygotes due to inbreeding in generally smaller, more isolated southern populations, but that this effect has not yet led to a loss of overall genetic diversity at the population level (*H*_*e*_, *π*).

We also note that far northern and far southern populations of both species sometimes exhibit slightly lower population genetic diversity (*H*_*e*_, *π*) than mid-latitude populations (Fig. 2). In this case, we do suspect that this pattern reflects reduced gene flow between core and peripheral populations and loss of diversity due to genetic drift in generally smaller, more isolated peripheral populations (Hampe & Petit, 2005; Jaramillo-Correa et al., 2009). However, in some European taxa, elevated mid-latitude genetic diversity is believed to instead reflect historical secondary contact and admixture of lineages from distinct southern refugia (Petit et al., 2003). Nonetheless, we find no evidence of distinct refugia or mid-latitude admixture among lineages, making secondary contact a less likely explanation for the patterns we observe.

### Temperate forests in ENA during the LGM

The phylogeographic history of both *C. cordiformis* and *C. ovata* is best characterized by simple latitudinal range shifts. Our genetic data and species distribution models suggest that both species inhabited large areas of the southeastern United States during the Last Glacial Maximum and gradually expanded northward to occupy their current distribution as glaciers retreated. We find no evidence of highly genetically distinct geographic regions or of strong phylogeographic breaks that would suggest population fragmentation into multiple refugia (except possibly range-edge *C. ovata* from Texas; see below). We also find no evidence that postglacial recolonization of northern areas has occurred via multiple routes or been impeded by major landscape barriers. LGM survival over large areas of the southeastern United States and gradual northward recolonization is also compatible with the fossil record for the genus *Carya* (Prentice et al., 1991; Jackson et al., 2000).

While we do not detect any signatures of complex refugial dynamics or multiple postglacial recolonization routes, an alternative scenario we must consider is that one or both species might have experienced a more complex phylogeographic history, the signatures of which have been subsequently obscured by extensive contemporary gene flow (e.g., He et al., 2013). Low population genetic differentiation and an IBD pattern suggest that gene flow among populations is indeed fairly high. However, both species are slow growing and long lived, with peak reproduction occurring in *C. cordiformis* from ages 50 to 125 (Smith, 1990), and in *C. ovata* from ages 60 to 200 (Graney, 1990). Relatively few generations have therefore passed since the LGM, meaning that there has likely been insufficient time for gene flow to erode genetic signatures of postglacial expansion from multiple refugia.

Given that *C. cordiformis* and *C. ovata* are common, widespread tree species with a geographic distribution roughly matching that of temperate forests in ENA, we consider them to be model taxa for understanding the phylogeographic history of temperate forests from this region as a whole. Both species occupy a variety of habitat sites, but do show relevant differences in their ecology (Graney, 1990; Smith, 1990). In particular, *C. ovata* is a habitat generalist, and phylogeographic patterns for this species are likely to be broadly representative of those for ENA forest taxa in general. In contrast, *C. cordiformis* is a more mesic, bottomland species in many parts of its range, and thus would be expected to be particularly susceptible to fragmentation into refugia and therefore exhibit strong phylogeographic structure. However, low genetic structure and lack of any evidence of distinct refugia in *C. cordiformis* provide particularly strong evidence that temperate forests in ENA during the LGM were not highly fragmented.

Several other widespread woody plant species from ENA are also believed to have expanded from a large area covering much of the southeastern United States, including *Acer rubrum* (McLachlan et al., 2005), *Dirca palustris* (Peterson & Graves, 2016), *Fagus grandifolia* (Bennett, 1985; McLachlan et al., 2005), and *Quercus rubra* (Magni et al., 2005). The presence of climatic conditions able to support temperate trees over large areas of the southeastern United States during the LGM is also well supported by pollen records (Prentice et al., 1991; Jackson et al., 2000; Williams, 2002). Despite these sources of phylogeographic concordance, more complex scenarios involving divisions into distinct refugia or multiple recolonization routes have been proposed for many other plant species, especially those that are less geographically widespread (e.g., Griffin & Barrett, 2004; Gonzales et al., 2008; Eckert et al., 2010; Barnard-Kubow et al., 2015; Nadeau et al., 2015; Zinck & Rajora, 2016). In addition, many forest communities with no modern analogue were present in ENA during the LGM (Jackson et al., 2000), suggesting that not all taxa responded to glaciation in the same way. It is therefore unclear whether any general migrational responses to glaciation are applicable to ENA forests, or whether most taxa exhibited species-specific, idiosyncratic responses.

One possible explanation for different responses among species is that suitable habitat for species adapted to a narrower range of climatic or edaphic conditions could have been more geographically fragmented during the LGM than for more widespread species. In particular, ENA taxa with a strictly southern, warm-temperate distribution may have become fragmented into distinct far-southern refugia in Florida, Texas, or along the Gulf or Atlantic Coasts (e.g., Gonzales et al., 2008; Eckert et al., 2010). In contrast, more widespread species such as *C. cordiformis* and *C. ovata* could have survived in more expansive inland areas of cool-temperate conditions that extended farther north (e.g., Fig. 5). As model taxa, *C. cordiformis* and *C. ovata* may be most appropriate for understanding the phylogeographic history of temperate forests as a whole, and may be less representative of taxa with widely differing habitat requirements and narrower ecological amplitudes.

Nonetheless, our results suggest that temperate forests from ENA may have experienced a relatively simple phylogeographic history. Previous phylogeographic studies have established that major rivers and the Appalachian Mountains are important phylogeographic barriers for some ENA taxa, but these findings are not universal and are typically found in small animals with limited dispersal ability (Soltis et al., 2006). In contrast, rivers are unlikely to present substantial dispersal barriers to large trees, and the phylogeographic impact (or lack thereof) of the Appalachians on tree populations remains unclear (Jaramillo-Correa et al., 2009). Overall, there are few topographic or climatic barriers likely to lead to complex phylogeographic histories for ENA forest trees, especially in comparison to other temperate regions of the world (Hewitt, 2000; Shafer et al., 2010; Qiu et al., 2011).

### Genetic distinctiveness of Texas shagbark hickory

Although an IBD pattern characterizes genetic structure throughout most of ENA in both species, the Texas population (TX) of *C. ovata* presents an exception to this general pattern. TX forms its own PCA genetic cluster that does not overlap with any other population (pink dots; Fig. 4b), and it is the only population with membership almost exclusively in the western cluster identified by FASTSTRUCTURE.

There are several possible interpretations of these results. One possibility is that TX may be derived from a separate glacial refugium. Glacial refugia have previously been inferred in Texas and northern Mexico for several southern *Pinus* and *Prunus* species (Schmidtling & Hipkins, 1998; Schmidtling, 2003; Shaw & Small, 2005; Jaramillo-Correa et al., 2009; Eckert et al., 2010). Alternatively, ancestors of the TX population may have experienced gene flow with *C. ovata* populations in the mountains of northern Mexico (Little, 1971). Although we have not included any high-elevation Mexican populations in our genetic analyses or paleodistribution models, we cannot exclude the possibility that these populations may have migrated to lower elevations and come into contact with other populations during the LGM or at another time during the Pleistocene.

However, we do not necessarily need to invoke a separate refugium or contact with Mexican populations to explain the genetic distinctiveness of the range-edge TX population, because our results could also merely reflect low gene flow between TX and other populations. There is likely very little gene flow between TX and populations we sampled further to the east due to the absence of *C. ovata* in the Lower Mississippi River Valley (Fig. 1). In addition, the Ozark and Ouachita Mountains could conceivably reduce gene flow with populations sampled to the north. Denser population sampling across the southwestern portion of the species range could help fill in geographic gaps in our PC analyses, in order to determine whether TX is truly a distinct gene pool, or represents the extreme end of a rangewide IBD pattern, but is subject to greater genetic isolation than other populations.

Given the uncertain origin of the Texas population of *C. ovata*, we suggest that any conservation efforts in this species should ensure inclusion of populations from Texas and surrounding regions. Genetically distinct southern populations of temperate species have often been identified as high conservation priority (Petit et al., 2003; Hampe & Petit, 2005; Médail & Diadema, 2009). However, across most of the range of *C. cordiformis* and *C. ovata*, we see little reason to prioritize conservation of southern populations, due to their lack of genetic distinctiveness and lack of elevated genetic diversity. On the other hand, most temperate tree species exhibit geographically structured climatically adaptive genetic variation (Savolainen et al., 2007; Aitken & Bemmels, 2016) unlikely to be captured by the putatively neutral SNP markers we employed. Conserving populations from a variety of climates across the species range would therefore likely maximize conservation of adaptively relevant genetic diversity.

## Conclusions

Genome-wide patterns of genetic variation, predictions of paleodistibution models, and the fossil record all suggest that *C. cordiformis* and *C. ovata* recolonized postglacial ENM from a single, continuous region of temperate forest that likely covered much of the southeastern United States during the LGM. We generally find no evidence of distinct glacial refugia or strong phylogeographic breaks. However, as an exception to this overall pattern, it is unclear whether the higher genetic distinctiveness of a Texas population of *C. ovata* relative to other populations reflects the effects of a separate Texas refugium, historical contact with Mexican populations, or contemporary patterns of gene flow. The two *Carya* species we have studied represent excellent model taxa for understanding the phylogeographic history of eastern deciduous forests as a whole. Whereas complex refugial dynamics must be invoked to explain genetic structure in other temperate regions of the world, our results suggest that the phylogeographic history of temperate forests from ENA can largely be explained by simple latitudinal range shifts, without division into distinct refugia.

## Acknowledgements

The authors thank B.J. Belcher, P. Cousineau, R. D’Andrea, J. Fiske, J. Ronson, D. Saenz and P. Tichenor for assistance with fieldwork, and A.T. Thomaz for assistance with genetic analyses. Graduate student support and research funding was provided to J.B.B. by the National Science Foundation (GRFP fellowship; DDIG 1501159), and the University of Michigan EEB Department and Rackham Graduate School.

## Data accessibility

SNP datasets and species occurrence records are available for download from the Dryad Digital Repository (DOI: XXXXX).

## Biosketches

Jordan Bemmels is interested in biogeography of temperate and tropical trees and this work is part of his PhD thesis. Christopher Dick is his thesis advisor, and is broadly interested in tropical tree biogeography and evolution.

## Author contributions

J.B.B. and C.W.D. conceived the project; J.B.B. performed fieldwork and labwork, and analyzed the data; J.B.B. wrote the manuscript with input from C.W.D.

